# Rhythmic Entrainment Echoes in Auditory Perception

**DOI:** 10.1101/2022.12.07.519456

**Authors:** Sylvain L’Hermite, Benedikt Zoefel

## Abstract

Rhythmic entrainment echoes – rhythmic brain responses that outlast rhythmic stimulation – can evidence endogenous neural oscillations entrained by the stimulus rhythm. We here tested for such echoes in auditory perception. Participants detected a pure tone target, presented at a variable delay after another pure tone that was rhythmically modulated in amplitude. In four experiments involving 154 participants, we tested (1) which stimulus rate produces the strongest entrainment echo and (2) – inspired by audition’s tonotopical organisation and findings in non-human primates – whether these are organized according to sound frequency. We found strongest entrainment echoes after 6-Hz and 8-Hz stimulation, respectively. Best moments for target detection (in or in anti-phase with the preceding rhythm) depended on whether sound frequencies of entraining and target stimuli matched, in line with a tonotopical organisation. However, for the same experimental condition, best moments were not always consistent across experiments. We provide a speculative explanation for these differences that relies on the notion that neural entrainment and repetition-related adaptation might exercise competing, opposite influences on perception. Together, we find rhythmic echoes in auditory perception that seem more complex than those predicted from initial theories of neural entrainment.

## Introduction

Rhythmic stimulation, both sensorily and electrically, produces rhythmic patterns in neural and perceptual measures that are synchronised with the rhythm of stimulation^1–3^. This effect is often called “neural entrainment”^4,5^ and assumed to involve neural oscillations, i.e. brain activity that is endogenously rhythmic^1^. This assumption is difficult to verify during stimulation, as any rhythmicity in neural or perceptual responses can be due to the rhythmicity of the stimulus, without involving endogenous neural oscillations^6,7^.

Rhythmic entrainment echoes are rhythmic brain responses that are produced by a rhythmic stimulus and persist after its offset. Endogenous oscillations should linger for some time after having been entrained, similar to a swing that has been pushed, whereas other, evoked brain activity will disappear rapidly when no stimulus is present^6,8^. Rhythmic entrainment echoes therefore play a crucial role in distinguishing entrained neural oscillations from other brain responses that are not endogenously rhythmic.

Recently, several studies have reported rhythmic entrainment echoes (sometimes called “forward entrainment”^9^). These have been observed after visual flicker^10^ and speech sounds^8,11^ in human neurophysiological recordings, after regular tone sequences in the auditory cortex of non-human primates^12^, and even after transcranial alternating current stimulation (tACS) in speech perception^8^.

We here examined rhythmic entrainment echoes in human auditory perception. We presented participants with a rhythmically amplitude-modulated (AM) pure tone, followed by a target tone that they were asked to detect (Fig. 1). We hypothesized that the AM tone would entrain oscillations and produce an entrainment echo. The target was presented at different delays relative to the AM tone and thus used to “probe” this echo.

**Figure 1.**
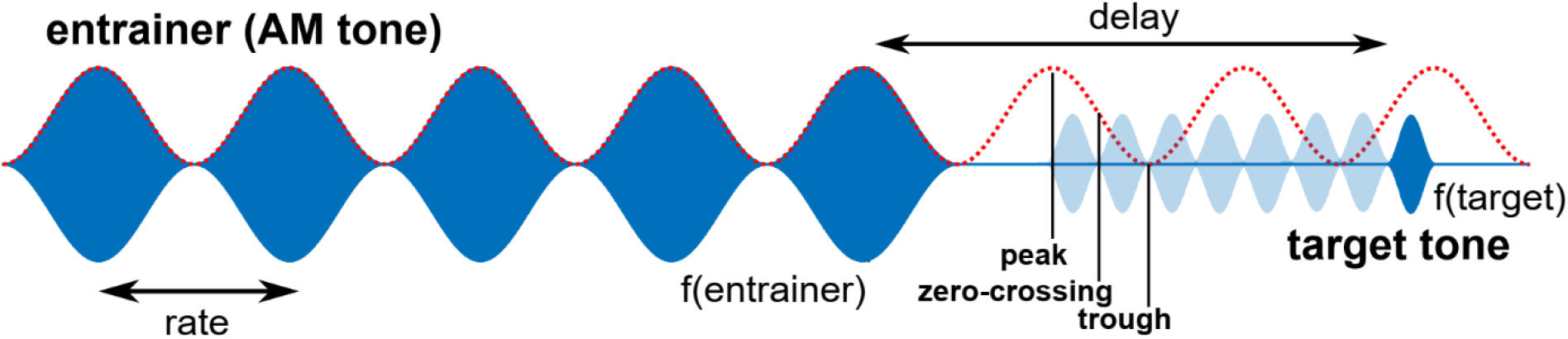
Experimental paradigm. A target pure tone with a sound frequency f(target) was presented at variable delays after a rhythmically amplitude-modulated (AM) tone with a sound frequency f(entrainer) and presented at a certain rate. We hypothesized that the AM tone would produce an echo in auditory perception that follows its rhythm (red dotted line). All possible target delays are shown for Experiment 1. One of them was selected randomly in each trial. In all experiments, targets were presented at the peak, zero-crossing, or trough of the preceding AM rhythm (i.e. four delays per cycle).

In a similar paradigm, Hickok et al.^13^ showed that AM noise, presented at 3 Hz, produces a 3-Hz oscillation in the detection of a subsequently presented a target tone. In that study, targets were most likely to be detected when presented in anti-phase with the preceding AM noise. We followed up on this finding, guided by two principal questions:

1. What is the preferred rate (*eigenfrequency*) of oscillations in audition? Stimulus rates that are closest to the “natural” frequency of neural oscillations should produce strongest entrained oscillations^14^ and, consequently, strongest entrainment echoes. A single previous study^15^ reported that these echoes are strongest for relatively slow rates (~2-3 Hz), but sample size was low (N = 3-5). We varied the rate of the AM tone and tested which of those rates produces the strongest oscillation in subsequent target detection.
2. Are entrainment echoes tonotopically organised? Previous work in non-human primates^12,16^ showed that rhythmically presented pure tones with a sound frequency *f* entrain neural activity in most of primary auditory cortex (A1). However, neuronal ensembles in parts of A1 that are tuned to (i.e. respond most strongly to) *f* aligned their high-excitability phase to the tones, whereas those in other parts aligned the opposite, low-excitability phase. This suggests that neural entrainment, and consequently entrainment echoes, are organised according to sound frequency. We independently varied the sound frequency of AM tone (*f_entrainer_*) and that of the target tone (*f_target_*) to test this hypothesis. More precisely, we hypothesized that the proportion of detected targets is highest in phase with the preceding AM tone if *f_entrainer_* = *f_target_*, but in anti-phase if *f_entrainer_* ≠ *f_target_* (Fig. 2A).

**Figure 2.**
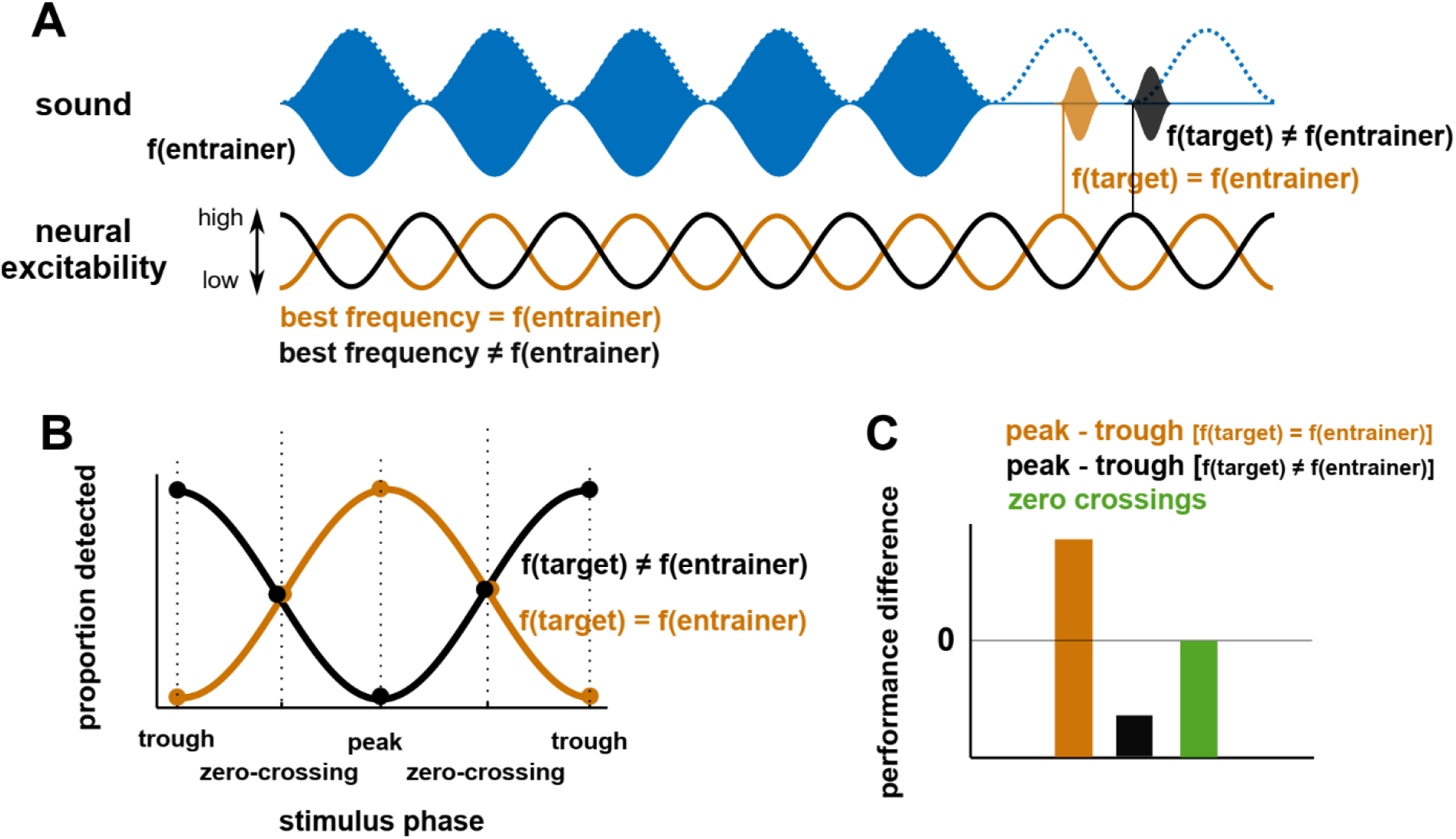
Frequency-specific entrainment echoes and their quantification. A. Based on findings in primary auditory cortex of non-human primates, we assumed that a rhythmically presented AM tone synchronises neural excitability “tonotopically”. That is, high-excitability phases only align to AM peaks if the sound frequency of the AM tone (f_entrainer_) matches the “best frequency” (i.e. the sound frequency that produces the strongest response) of the region the oscillation is measured in (orange); otherwise the opposite, low-excitability phase aligns (black). Note that the role of the target tone is to “sample” such frequency-specific oscillations: The probability of detecting this target tone should follow excitability fluctuations in the region processing the sound frequency of that tone (f_target_). B,C. For f_entrainer_ = f_target_, our hypothesis predicts entrainment echoes in phase with the AM rhythm, reflected by (B) the highest proportion of detected targets at the peak of the preceding AM rhythm, and (C) a positive difference for peak vs trough performance. For f_entrainer_ ≠ f_target_, it predicts entrainment echoes in anti-phase with the AM rhythm, reflected by (B) a highest proportion of detected targets at the trough of the AM tone, and (C) a negative difference for peak vs trough performance. In both cases, it predicts a peak-trough performance difference that is larger than that between the two zero-crossings tested (green in C).

## Methods

### Participants

We tested for entrainment echoes in four independent experiments. Experiment 1 was run in the laboratory. 16 participants (9 female; mean 26.2 years, range 23-34 years) completed the experiment after giving written informed consent.

Experiments 2-4 were run online, in chronological order as they are described here. 55, 59, and 50 participants were recruited from Prolific Academic (www.prolific.co, cf. ref ^17^) for those three experiments and gave informed consent by clicking on a button to confirm they wanted to participate. 8, 10, and 8 participants were excluded, either for failing a test designed to ensure they were wearing headphones, or because they did not respond accurately during practice trials (see Procedure). Consequently, data from 47 (10 female; mean 29.1 years, range 19-46 years), 49 (22 female; mean 32.8 years, range 19-46 years), and 42 (15 female; mean 28.7 years, range 19-44 years) participants were included for subsequent analyses for Experiments 2, 3, and 4, respectively.

This study was approved by the Comité de Protection des Personnes (CPP) Ouest II Angers (protocol number 2021-A00131-40).

### Stimuli

In all experiments, participants were presented with AM pure tones at a certain rate, followed by a target tone that they were asked to detect. The target tone was present in 50% (Experiment 1) or 33.33 % (Experiments 2-4) of the trials and its level was adapted to an individual threshold (see Procedure). The duration of AM and target tones was always 5 and 0.25 cycles of the presentation rate, respectively. The delay between the two was defined as the onset of the target tone relative to the final peak of the AM tone (Fig. 1). This delay was variable and the critical experimental manipulation to reveal entrainment echoes: If such echoes existed, then probability of target detection should depend on the time of target presentation relative to the offset of the AM tone. Several acoustic parameters differed between experiments and are summarised in Table 1 (and shown in Fig. 1). These are the rate of the AM tone, the sound frequencies of both AM tone (*f_entrainer_*) and target (*f_target_*), and the exact delays between AM and target tones. The step size between possible delays was 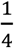 cycle of the AM rhythm in all experiments so that targets were presented at the peak, trough, or the two zero-crossings of the preceding AM stimulus (Fig. 1).

In Experiment 1, we tested which of five different rates produces the strongest entrainment echo. We only used a single combination of sound frequencies (*f_entrainer_* = 500 *Hz, f_target_* = 1200 *Hz*) to increase the number of trials per condition. This experiment revealed echoes that are strongest after 6-Hz AM tones and around 2-3 cycles after entrainer offset.

For Experiment 2, we therefore restricted the rate of AM tones to 6 Hz and used longer delays. We used different combinations of sound frequencies (divided into the two categories *f_entrainer_ = f_target_* and *f_entrainer_* ≠ *f_target_*) to test the hypothesis that entrainment echoes are tonotopically organised (see Introduction and Statistical Analysis).

Whereas *f_entrainer_* and *f_target_* were identical (and therefore predictable) in each trial in Experiment 1, they varied (unpredictably) from trial to trial in Experiment 2. In Experiment 3, we tested whether differences in outcomes from the two experiments (see Results) are due to these differences in the predictability of *f_target_*. This was done by varying the predictability of *f_target_* in different experimental blocks (see Procedure). We also adapted the delays between AM tone and target tone to those that showed strongest effects in Experiments 1 and 2.

**Table 1.**
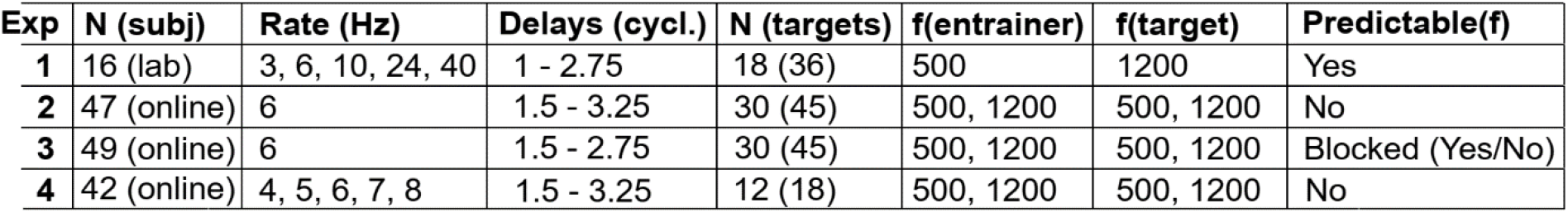
Summary of experimental parameters used in the different experiments. Delays are expressed as the number of cycles the target tone was presented at after the final peak of the AM tone. In all experiments, the step size for delays was 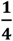 cycle. Consequently, 8 delays were tested in Experiments 1, 2, and 4, and 6 delays in Experiment 3. Targets/delay is the number of target trials per condition (e.g., rate) and delay. The total number of trials per condition and delay is shown separately in brackets. Predictability refers to the fact that participants were able to predict the sound frequencies of AM tone and target tone in Experiment 1 (as only a single combination was used), but not in Experiments 2 and 4 (as they were randomly selected in each trial). In Experiment 3, predictability of sound frequency was varied block-wise (see Procedure).

| Exp | N (subj) | Rate (Hz) | Delays (cycl.) | N (targets) | f(entrainer) | f(target) | Predictable(f) |
| --- | --- | --- | --- | --- | --- | --- | --- |
| 1 | 16 (lab) | 3, 6, 10, 24, 40 | 1 - 2.75 | 18 (36) | 500 | 1200 | Yes |
| 2 | 47 (online) | 6 | 1.5 - 3.25 | 30 (45) | 500, 1200 | 500, 1200 | No |
| 3 | 49 (online) | 6 | 1.5 - 2.75 | 30 (45) | 500, 1200 | 500, 1200 | Blocked (Yes/No) |
| 4 | 42 (online) | 4, 5, 6, 7, 8 | 1.5 - 3.25 | 12 (18) | 500, 1200 | 500, 1200 | No |

In Experiment 1, our sampling of rate for the AM tone was relatively sparse (see Table 1). In Experiment 4, we again varied this rate, but in a narrower range, centred around the one that produced the strongest echo in Experiment 1 (6 Hz). This manipulation had several purposes: It allowed us (1) to estimate the “preferred” rate for entrainment echoes with a finer spatial resolution; (2) to vary rate and sound frequencies (*f_entrainer_* and *f_target_*) within the same experiment; and (3) to test whether differences in outcomes between Experiment 1 and Experiments 2 and 3 (see Results) are due to rate being variable across trials only in the former.

### Procedure

Participants’ task was fairly simple and similar across experiments: They were asked to indicate with a button press whether they perceived a target tone. In Experiments 1 and 3, this corresponded to a simple yes/no (forced) choice. In Experiments 2 and 4, they had to choose between high-pitch, low-pitch, and no target present. This was done to obtain false alarms that can be interpreted more easily, but not feasible in the other experiments as these included experimental blocks with only one possible *f_target_* (which made the high vs low-pitch differentiation meaningless).

Experiments 2-4 were conducted over the internet using the jsPsych JavaScript library^18^ and Cognition experiment management software. Mobile phones and tablets were ineligible, which was ensured with a JavaScript-based device check. Participants were asked to complete the experiment in a quiet, room, and to wear headphones.

All experiments began with a calibration sound (pure tone) to verify that the audio could be heard clearly, and to allow participants to adjust their volume to a comfortable level. Participants were instructed not to adjust the volume on their computer for the remainder of the experiment.

For online experiments only, this was followed by a test designed by Woods et al.^19^ to ensure that participants were wearing headphones. This test can be easily completed when wearing headphones but not otherwise. A detailed description of can be found in ref ^20^.

In all experiments, participants then received detailed instructions about the task and listened to example sounds. They were asked to complete practice trials in which the target was clearly audible. Participants received feedback on whether they responded correctly after each practice trial. In online experiments, participants were only able to continue with the main part of the experiment if they responded correctly in at least 4 out of 5 practice trials. In case of failure, they were allowed to repeat practice once. In the lab experiment, all participants completed practice successfully.

Subsequently, the level of the target tone was adjusted to individual participants’ detection thresholds. Participants were presented with the same stimuli that were used for the main experiment, with a randomly chosen delay between AM tone and target tone in each trial. The level of the target tone decreased or increased in each trial. Participants were instructed to press a button when they could no longer hear the tone (for the decreasing level sequences) and when they started to hear it again (for the increasing level sequences). Decreasing and increasing level sequences were used in alternation. Participants’ detection threshold was defined as the average level of the target tone during the last four button presses. This adaptation procedure was run separately for each rate (Experiments 1 and 4) and combination of *f_entrainer_* and *f_target_* (Experiments 2-4).

Finally, participants completed the main experimental task, as described above. In each trial, (1) the level of the target tone (−2 dB, 0 dB or +2 dB relative to the individual threshold; in Experiment 4, only 0 dB was used), (2) the delay between AM tone and target tone, (3) the rate of the AM tone (only in Experiments 1 and 4), and (4) the combination of *f_entrainer_* and *f_target_* (only in Experiments 2-4) were selected pseudo-randomly. This part was divided into several experimental blocks. In Experiment 3, there were two types of blocks. In some blocks, the combination of *f_entrainer_* and *f_target_* was identical in each trial and therefore predictable. In other blocks, it was selected pseudo-randomly in each trial (out of four possibilities) and therefore unpredictable. Prior to each block, participants were informed about block type (including which combination of sound frequencies will be presented), selected pseudo-randomly. Between blocks, participants were offered a break and told that they could continue with the task by pressing a key when they were ready.

### Statistical Analysis

For each participant, delay, and experimental condition (e.g., AM rate), we computed the proportion of correctly detected targets (*p_hi_t*). Rhythmic entrainment echoes would be visible as rhythmic fluctuations in *p_hit_* after the offset of the AM tone, and at the corresponding rate.

We tested for entrainment echoes in “sliding windows” that each comprised four delays (i.e. one full cycle), and using a step size of one delay (i.e. 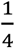 cycle). These four delays corresponded to the peak, trough, and the two zero-crossings of the preceding AM rhythm (Fig. 1). This allowed us to estimate entrainment echoes with a finer temporal resolution as compared to an approach that combines all delays tested into a single estimate.

Our hypothesis makes clear predictions about best and worst moments for target detection, shown in Fig. 2B. In particular, for *f_entrainer_ = f_target_*, we expected *p_hit_* to be highest and lowest in phase and in anti-phase with the AM rhythm (i.e. at its peak and trough, had it continued), respectively, whereas the opposite should be true if *f_entrainer_ ≠ f_target_*. On the group level, we tested this prediction by computing the corresponding difference in performance (peak – trough) for each participant and by comparing the outcomes against the null hypothesis of 0 (t-test), or across conditions (e.g., rate; repeated-measures ANOVA).

We used two additional analytical steps to rule out other factors that could have produced the expected peak-trough performance difference without the cyclic pattern of performance (Fig. 2B) that characterises entrainment echoes. First, we corrected for linear changes in detection performance over time by removing a linear fit from each cycle before computing performance differences described above. Second, we used the fact that – apart from the positive or negative difference in performance at peak and trough (orange and black in Fig. 2C) – our hypothesis predicts a near-zero difference between the two zero-crossings (green in Fig. 2C). We tested for this by contrasting (via paired t-tests) the two corresponding differences (peak-trough vs zero-crossings). As performance should be similar for both zero-crossings, the order of the two performance values (first minus second zero-crossing or vice versa) was chosen randomly.

We here note that our assumption – highest *p_hit_* in phase or in anti-phase with the preceding AM rhythm (but not in-between) – relies critically on the use of *perceptual* outcomes as a measure of entrainment echoes. This assumption is not necessarily true for neurophysiological measures of entrainment echoes, due to individual delays between the presentation of a stimulus and its effect on those neural measures^8^. Such delays, however, should be *identical* for AM tone and target tone and therefore not affect perceptual outcomes.

For each participant and condition, we also determined the proportion of false alarms (*p_false alarm_*), i.e. target-absent trials that were incorrectly labelled as “target present”. In Experiments 2 and 4, we computed *p_false alarm_* separately for the two possible target sound frequencies. We used false alarms to compute participants’ sensitivity to targets, quantified as:

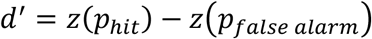

Note that *p_hit_* is defined for individual delays, but *p_false alarm_* is not, because false alarms are not defined for specific delays (as no target is present). For each d’, the same *p_false alarm_* (but a different *p_hit_*) was therefore used. Despite *p_false alarm_* being constant, a recent study^21^ showed that this approach can still produce different results when comparing entrainment echoes in proportion correct (hits) and sensitivity (d’). Using the approach described above, we thus tested whether entrainment echoes can be observed in d’.

## Results

In four experiments, we tested whether the detection of a target tone fluctuates rhythmically at the rate of a preceding amplitude-modulated (AM) tone (Fig. 1), revealing entrainment echoes in auditory perception.

The simplest version of the “neural entrainment theory” predicts best perception in phase with a rhythmic stimulus, as high-excitability moments of the oscillatory cycle synchronise with the expected timing of upcoming events^1,22^. Studies on non-human primates in primary auditory cortex^12,16^ confirmed this assumption when neural activity was measured in areas “tuned” to the sound frequency of the entraining (rhythmic) stimulus. However, the opposite effect seems to occur (high excitability in anti-phase with the rhythmic stimulus) for other sound frequencies (Fig. 2A).

We quantified rhythmic entrainment echoes by testing two predictions that follow from these previous findings, and from the expected cyclic shape of the perceptual modulation (Fig. 2B). We first computed proportions of detected targets that were presented in phase and in anti-phase with the preceding AM tone, respectively (i.e. at its peak vs trough, had it continued). The difference between in and anti-phase target detection (*peak-trough difference*) should be different from 0 (Fig. 2C), whereas the sign of the difference reflects the phase of the echo (positive and negative for in phase and anti-phase entrainment echoes, respectively). We then compared this difference (peak-trough; orange and black in Fig. 2C) with the corresponding difference between two zero-crossings of the putative rhythmic echo, and that we predicted to be near-zero (green in Fig. 2C).

In a first lab experiment, we tested which stimulus rate produces the strongest entrainment echoes. In three follow-up online experiments, we tested the hypothesis that entrainment echoes are frequency-specific (Fig. 2), leading to simultaneous best and worst moments for the detection of a target, depending on whether its sound frequency differs from that of the entraining stimulus (AM tone), respectively. In Experiment 4, we also estimated the “preferred” stimulus rate for entrainment echoes with a finer spatial resolution.

### Experiment 1 (N = 16, lab experiment)

In Experiment 1, we varied the rate of the entraining tone’s amplitude modulation. The same sound frequencies were used in each trial, and differed between AM tone (*f_entrainer_* = 500 *Hz*) and target tone (*f_target_* = 1200 *Hz*).

On average, participants detected 50.0 % (± 13.1 %; SD) of the targets and made 4.6 % (± 3.4 %) false alarms (resulting in an average d-prime of 1.88 ± 0.37). There was no difference in the proportion of detected targets across rates (F(4) = 1.24, p = 0.30; repeated-measures ANOVA), indicating that target levels were successfully adapted to individual detection thresholds for all rates (see Procedure). There was, however, a difference in false alarm probability (F(4) = 3.04, p = 0.02), with fewer false alarms after 10-Hz AM tones than after 24-Hz and 40-Hz AM tones (all other post-hoc comparisons were non-significant). D-prime measures did not differ across rates (F(4) = 0.52, p = 0.72).

For each rate, we then tested for rhythmic entrainment echoes in different temporal windows – each one cycle long – after the offset of the AM tone (see Statistical Analysis). We found an entrainment echo, exclusively after 6-Hz stimulation (Fig. 3A) and in the last cycle tested (continuous line in Fig. 3B). This echo was illustrated by a negative difference between peak and trough performance that was reliably different from 0 (t(15) = 4.43, FDR-corrected p = 0.006, effect size Cohen’s d = 1.11). This peak-trough difference (black in Fig. 3A) was also significantly different (t(15) = 4.82, FDR-corrected p = 0.006, Cohen’s d = 1.20) from the difference between zero-crossings (blue in Fig. 3A), as predicted from a cyclic pattern in perceptual outcomes (see Statistical Analysis and Fig. 2C). Importantly, the highest number of targets were detected in anti-phase with the preceding AM tone (Fig. 3B). This is predicted from tonotopic entrainment, as *f_entrainer_* and *f_target_* differed in this experiment (Fig. 2).

**Figure 3.**
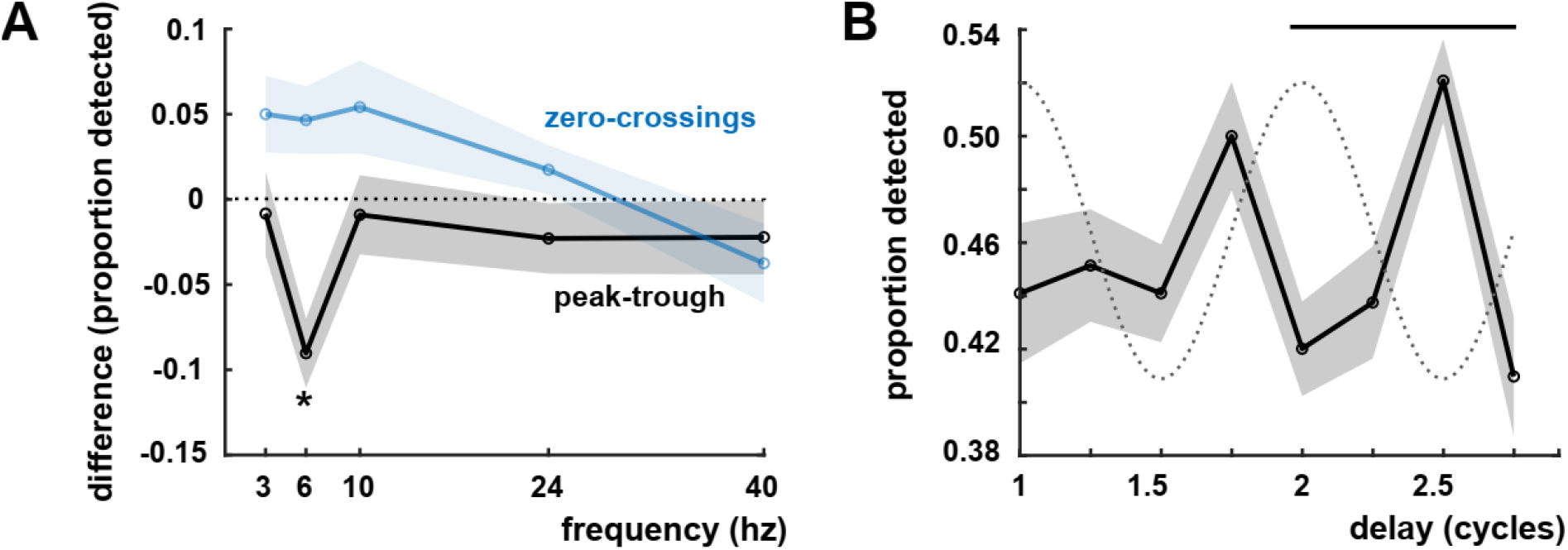
Experiment 1 Results. A. Difference in the proportion of detected targets, presented in phase (peak) and in anti-phase (trough) with the preceding AM rhythm (black), or at the two zero-crossings (blue). The negative sign reflects an anti-phase echo (cf. Fig. 2B,C). Results are shown only for the last cycle tested. B. Proportion of detected targets after the 6-Hz AM stimulus (results for all rates are shown in A). The dashed line shows the rhythm of the preceding AM tone, had it continued. The continuous line shows the temporal windows in which the peak-trough difference in detected targets (black in A) differs reliably from 0 and from the corresponding difference between zero-crossings (blue in A) (FDR-corrected p < 0.05). Shaded areas show standard error of the mean (SEM).

This result was confirmed by contrasting differences in peak-trough performance across conditions (black in Fig. 3A). Most importantly, we found an interaction of rate and delay (F(16) = 2.22, p = 0.005), reflecting echoes that are only present for certain rates (6 Hz) and for later delays tested.

Performance measures like d’ are a more complete indicator of participants’ perceptual sensitivity, as they combine the proportion of correctly detected targets (hits) with that of target-absent trials that were incorrectly labelled as “target present” (false alarms). In the current paradigm, however, false alarms are not defined for individual delays, as no target is present. Nevertheless, a recent study^21^ showed that entrainment echoes in hits and d’ can differ, even if the same (average) proportion of false alarms is used for each delay. Results for entrainment echoes in d’ are shown in Fig. S1A. We found very similar results as for hits, with an anti-phase entrainment echo only after 6-Hz stimulation and in later delays tested (FDR-corrected p-values < 0.01).

### Experiment 2 (N = 47, online experiment)

Based on results from Experiment 1, we fixed the rate of the AM tone to 6 Hz. As the entrainment echo was most apparent in the second cycle tested (Fig. 3B), we delayed possible presentation times of the target by half a cycle (see Table 1). The sound frequency of the AM tone (*f_entrainer_*) and that of the target tone (*f_target_*) could either be identical (condition *f_same_*) or different (condition *f_different_*), selected pseudo-randomly in each trial.

On average, participants detected 61.4 % (± 14.0 %) of the targets and made 1.6 % (± 2.5 %) false alarms (resulting in an average d-prime of 2.81 ± 0.66). There was no significant difference in target detection across conditions (*f_same_* vs *f_different_*; F(1) = 2.51, p = 0.12). However, when participants made a false alarm, they very frequently reported a target with sound frequency that is different from that of the AM tone (F(1) = 24.55, p < 0.0001), leading to a higher d’ for *f_same_* (F(1) = 63.28, p < 0.0001).

Fig. 4A shows how target detection fluctuated after the offset of the rhythmic AM stimulus, separately for *f_same_* (orange) and *f_different_* (black). Whereas most targets were detected in phase with the preceding rhythm for *f_different_*, maximal detection was observed in anti-phase for *f_same_*· In the corresponding temporal windows (continuous lines in Fig. 4A), these entrainment echoes were visible as peak-trough differences in detection that were larger than for zero-crossings (Fig. 4B). These differences reached or approached conventional statistical significance (*f_different_*, peak-trough vs 0: t(46) = 2.18, p = 0.03, Cohen’s d = 0.32; peak-trough vs zero-crossings: t(46) = 2.12, p = 0.04, Cohen’s d = 0.31; *f_same_*, peak-trough vs 0: t(46) = 2.68, p = 0.01, Cohen’s d = 0.39; peak-trough vs zero-crossings: t(46) = 1.94, p = 0.06, Cohen’s d = 0.28), but they did not survive correction for multiple comparisons (FDR-corrected p > 0.05).

**Figure 4.**
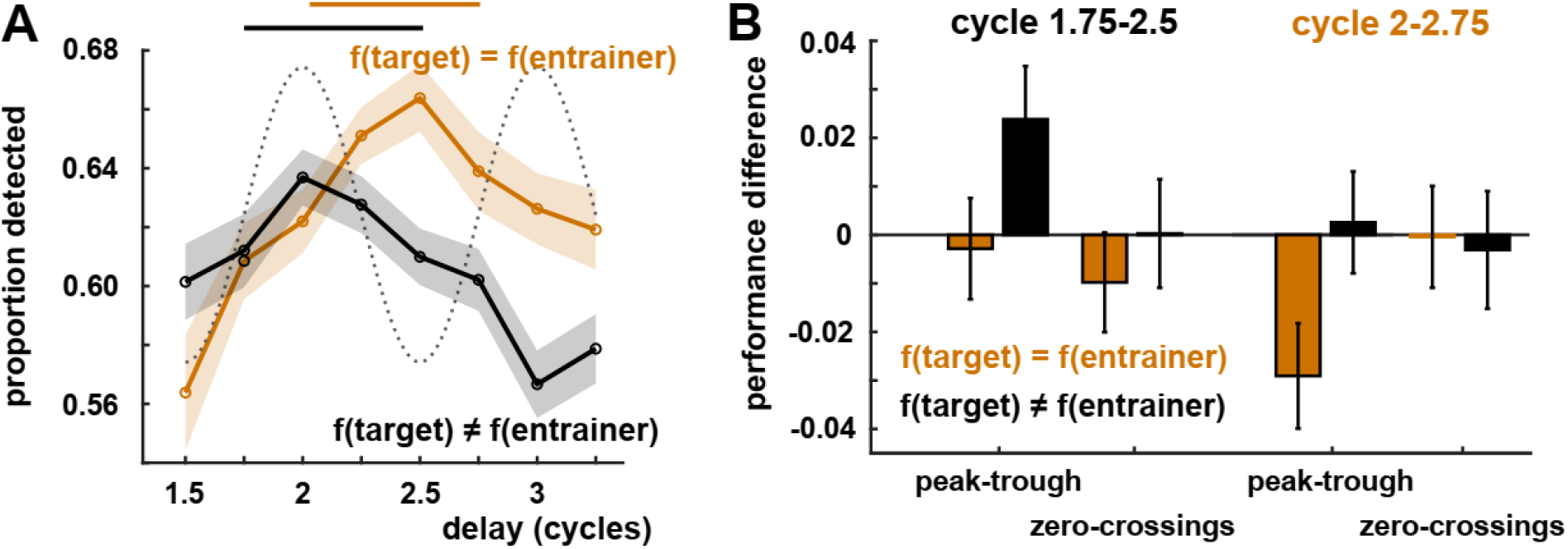
Experiment 2 Results. A. Proportion of detected targets as a function of target delay relative to the AM stimulus. Same conventions as for Fig. 3B. The lines show the temporal windows with strongest entrainment echoes, although these reach conventional statistical significance (p < 0.05) only if uncorrected for multiple comparisons. Note the opposite best AM phase for target detection in the f_different_ condition as compared to Experiment 1 (Fig. 3B). B. Differences in proportion of detected targets for peak-trough and the two zero-crossings in the respective cycles (cf. Fig. 2C). Results are only shown for the two cycles with the strongest entrainment echoes in the two conditions (continuous lines in A). Error bars show SEM.

An ANOVA on peak-trough differences revealed a main effect of temporal window (F(4) = 2.93, p = 0.02), driven by a change in maximal detection from in-phase to anti-phase (compare the two temporal windows shown in Fig. 4B). However, there was no main effect of condition (*f_same_* vs *f_different_*) nor interaction (p > 0.10). Results were again similar for d’ (Fig. S1B).

It is of note that (1) maximal detection for *f_different_* (in phase with preceding AM tone) is opposite to that observed in Experiment 1 (in anti-phase; note that *f_same_* was only tested in Experiment 2), and (2) the delays with strongest peak-trough differences are similar for the two experiments (between two and three cycles post-entrainer). This suggests that, despite their relatively small size, effects obtained in Experiment 2, and their difference to Experiment 1, are meaningful, an assumption that led to the experiment described next.

### Experiment 3 (N = 49, online experiment)

Experiments 1 and 2 differed in the predictability of sound frequencies: *f_entrainer_* and *f_target_* were constant and predictable in Experiment 1, but randomly selected and unpredictable in Experiment 2. In Experiment 3, we tested whether this difference can explain opposite phases of maximal target detection for *f_different_* (in anti-phase and in phase with the preceding AM tone, respectively). As in Experiment 2, we varied *f_entrainer_* and *f_target_*, but in some blocks these were randomly chosen (condition *f_unpredictable_*; replicating Experiment 2), and in other blocks they remained constant throughout the block (condition *f_predictable_*). Prior to each block, participants were informed of the block type (and of sound frequencies if *f_predictable_*). Overall performance followed a similar pattern to that observed in Experiment 2. Participants detected 63.0 % ± 14.5 % of the targets, made 3.0 % ± 3.2 % false alarms, leading to a d’ of 2.51 ± 0.67. Conditions did not differ in the proportion of detected targets (*f_predictable_* vs *f_unpredictable_*, p = 0.50; *f_same_* vs *f_different_*, p = 0.43). False alarms were lower when sound frequencies were predictable (*f_predictable_*; t(48) = 2.72, p = 0.009), although the resulting d’ did not significantly differ from the one in the unpredictable condition (t(48) = 1.25, p = 0.22).

Fig. 5A,B shows main results from Experiment 3, reminiscent of those obtained in Experiment 2 (Fig. 4). Again, the proportion of detected targets peaked approximately in phase with the preceding AM tone for *f_different_*, and in anti-phase for *f_same_*. This pattern was more obvious when sound frequencies were predictable (*f_predictable_*; Fig. 5B) than when they were not (*f_unpredictable_*; Fig. 5A). For *f_predictable_*, this was evidenced by peak-trough differences that differed reliably from 0 and, albeit less reliably, from the difference between zero-crossings. This was the case in the first four delays tested for *f_different_* (peak-trough vs 0: t(48) = 3.25, FDR-corrected p = 0.01, Cohen’s d = 0.46; peak-trough vs zero-crossings: t(48) = 2.15, FDR-corrected p > 0.05, Cohen’s d = 0.32) and in the last four delays for *f_same_* (peak-trough vs 0: t(48) = 4.26, FDR-corrected p = 0.001, Cohen’s d = 0.60; peak-trough vs zero-crossings: t(48) = 2.86, FDR-corrected p > 0.05, Cohen’s d = 0.41), see continuous lines in Fig. 5B. In addition, an ANOVA revealed an interaction between predictability and temporal window (F(96) = 3.33, p = 0.04). Post-hoc tests indicated that this was driven by stronger echoes for *f_predictable_* at later delays only. Fig. S1C,D show corresponding results for d’.

**Figure 5.**
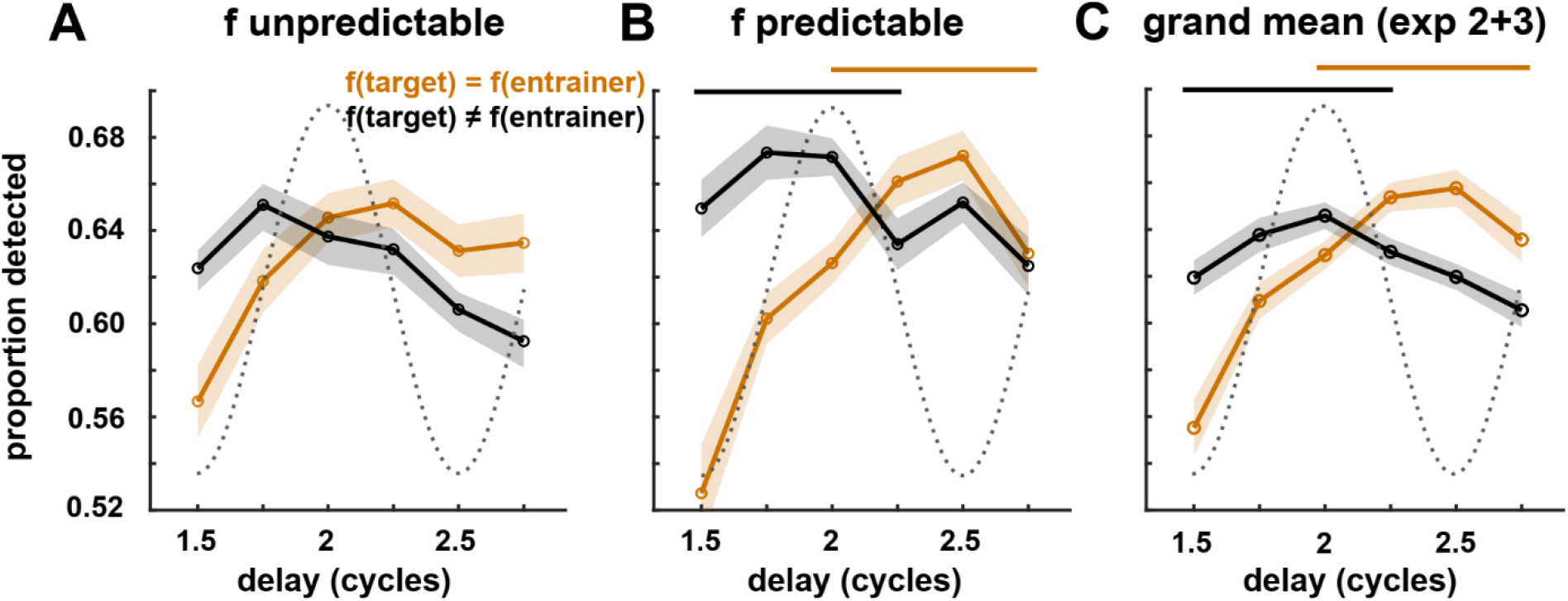
Experiment 3 Results. Same conventions as for Figs. 3B and 4A. The lines show the temporal windows with strongest peak-trough differences in the respective conditions (FDR-corrected p < 0.05). Panel C shows results for pooled subjects from Experiments 2 and 3.

Despite some statistically reliable results, entrainment echoes were relatively weak in Experiments 2 and 3. Given similar results in the two experiments, we therefore pooled their subjects (ignoring the differences in predictability of sound frequencies for this analysis). Results are shown in Fig. 5C, confirming peaks in the proportion of detection targets when they were presented in phase (*f_different_*, peak-trough vs 0: t(95) = 2.63, FDR-corrected p = 0.03, Cohen’s d = 0.27) and in anti-phase with the entraining stimulus (*f_same_*, peak-trough vs 0: t(95) = 3.65, FDR-corrected p = 0.003, Cohen’s d = 0.37), respectively. However, peak-trough differences were not reliably different from those between zero-crossings when correcting for multiple comparisons (all FDR-corrected p > 0.05, Cohen’s d = 0.17-0.24).

### Experiment 4 (N = 42, online experiment)

Results from Experiment 3 showed that a difference in the predictability of sound frequency cannot explain opposite phases of entrainment echoes in Experiments 1 and Experiments 2/3. In Experiment 4, we tested an alternative explanation for this effect. A repeated presentation of a given stimulus, such as pure tones used here, leads to progressive reduction of neural responses to this stimulus if it occurs at the expected time^23–25^, whereas any deviance in time or identity (e.g., sound frequency) produces a stronger response and possibly enhanced detection^26,27^ (for details, see Discussion). In our case, an enhanced detection of unexpected events would be visible as the highest proportion in detected targets in anti-phase with the preceding rhythm for *f_same_*, and in phase for *f_different_* – precisely what we observed in Experiments 2 and 3. This is a scenario that is *opposite* to that predicted by neural entrainment, where a rhythmic tone sequence prepares neural resources for the expected timing and identity of upcoming information, leading to optimal perception when a tone with the same sound frequency is presented in phase with the preceding rhythm. Our findings in Experiment 1 are in line with this prediction.

Importantly, in Experiment 1, but not in Experiments 2 and 3, the rate of the entraining AM tone varied across trials. It is possible that changes in rate prevented a repetition-related suppression of responses, leading to perceptual outcomes that are instead dominated by entrainment effects. We tested this hypothesis in Experiment 4. Again, we varied AM rate across single trials. Instead of using a wide range of rates as for Experiment 1, we used smaller steps around the rate that turned out to be optimal in Experiment 1 (4-8 Hz, in steps of 1 Hz). This also allowed us to test for “preferred” rates for entrainment echoes with a higher resolution. Participants detected 58.3 % ± 14.6 % of the targets, made 1.0 % ± 1.1 % false alarms, leading to a d’ of 2.70 ± 0.48. There was a main effect of rate on target detection (F(4) = 3.92, p = 0.005, repeated-measures ANOVA) that was driven by more detected targets after 4-Hz and 5-Hz AM tones (59.7 % and 59.6 %, respectively) than after 8-Hz tones (56.6 %). There was also a main effect of condition (F(1) = 14.90, p < 0.001), with more targets detected for *f_same_* (59.6 %) than for *f_different_* (57.1%). Moreover, we found a main effect of condition (but no main effect of rate) on the proportion of false alarms (F(1) = 36.24, p < 0.001), with more false alarms for *f_different_* (1.6 %) than for *f_same_* (0.5 %); and a main effect of rate (F(4) = 3.28, p = 0.01) as well as condition (F(1) = 81.39, p < 0.001) on d’, with a higher d’ for targets presented after 4 Hz (2.77) than 8 Hz AM tones (2.64), and a higher d’ for *f_same_* (2.90) than for *f_different_* (2.50).

Fig. 6 shows main results from Experiment 4. We again found an entrainment echo, this time exclusively after 8-Hz stimulation and for *f_same_* (Fig. 6A). This echo was most evident for earlier delays tested (continuous line in Fig. 6B) and confirmed by a statistically reliable difference in the detection of targets presented in phase and in anti-phase with the preceding AM tone (t(41) = 4.71, FDR-corrected p = 0.001, Cohen’s d = 0.73), respectively. This peak-trough difference also differed from the corresponding difference between zero-crossings (t(41) = 3.58, FDR-corrected p = 0.04, Cohen’s d = 0.55). As illustrated from the positive peak-trough difference (Fig. 6A), target detection was maximal in phase with the preceding AM rhythm, as predicted for *f_same_* from the neural entrainment hypothesis. However, for *f_different_* we did not find the corresponding anti-phase echo (cf. Fig. 2B) that was previously observed in Experiment 1 (Fig. 3B). Our results were confirmed by contrasting differences in peak-trough performance (black in Fig. 6A) across conditions. As for Experiment 1, we found an interaction of rate and delay (F(16) = 2.03, p = 0.01), reflecting echoes that are only present for certain rates (8 Hz) and for earlier delays tested. Fig. S1E shows results for d’, corroborating those described for the proportion of detected targets.

**Figure 6.**
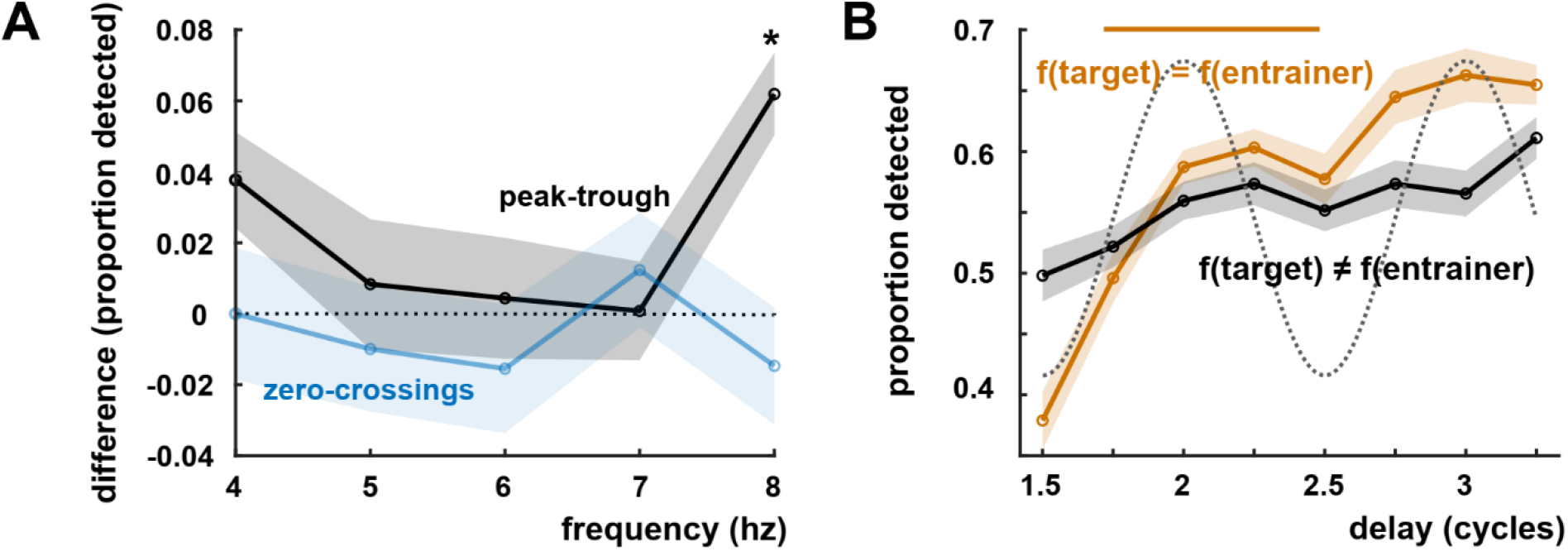
Experiment 4 Results. Same conventions as for Fig. 3. A shows differences between proportions of detected targets presented at peak and trough (black) and the two zero-crossings (blue) of the preceding AM rhythm, respectively. Results are shown only for the f(same) and the temporal windows with the strongest entrainment echo (continuous orange line in B). B shows how detection fluctuated in time, separately for the two experimental conditions and after the 8-Hz AM stimulus.

## Discussion

### Summary: Rhythmic entrainment echoes in auditory perception

Rhythmic brain responses that outlast rhythmic stimulation – “rhythmic entrainment echoes”^8,28^ or “forward entrainment”^9^ – do not only play an important role to demonstrate the involvement of endogenous brain oscillations^6,8^; they can also give us insights into whether and how participants predict the timing of events^9^, a fundamental notion for the fields of “neural entrainment”^4,5^ and “temporal attending”^29,30^.

Here, in four independent experiments, we examined entrainment echoes in auditory perception. Specifically, we asked (1) which stimulus rate leads to strongest echoes in the detection of a subsequent auditory target and (2) whether these effects are organised tonotopically (Fig. 2). For the latter, we hypothesised that pure tone targets are detected best if they are presented in phase with a preceding entraining stimulus at the same sound frequency, whereas detection is most likely in anti-phase when sound frequencies of target and entrainer differ. Notably, some previous studies reported peaks in performance or neural activity in phase with a preceding rhythmic stimulus^31,32^, whereas for others they occurred in anti-phase^10,13^. These results might support our hypothesis, but it remained to be tested whether sound frequency is indeed a decisive factor for these seemly opposing findings.

Indeed, not only did we find fluctuations in target detection that depended on the rhythm of the preceding AM stimulus – supporting the existence of entrainment echoes – these also changed their phase depending on whether sound frequencies of target and entraining stimulus matched. Surprisingly, however, for the same experimental condition (*f_sa_me* or *f_different_*), detection was maximal in phase only in some experiments, whereas it peaked in anti-phase in others (e.g., compare Figs. 3B and 4A for *f_different_*, and Figs. 4A and 6B for *f_same_*). This observation suggests that entrainment echoes are a phenomenon that is more complex that what seems to be predicted from initial theories of neural entrainment and dynamic attending.

### Entrainment vs habituation – two opposing processes?

Although the following does entail some speculation, differences between the four experiments conducted here can provide us with some clues about the origins of these seemingly opposite patterns in auditory perception. Such opposite findings (e.g., in phase vs in anti-phase echoes) were most prominent between Experiments 1 and 2/3 (anti-phase vs in phase echo for *f_different_*) and between Experiments 2/3 and 4 (anti-phase vs in phase echo for *f_same_*). We here propose that the observed differences might be due to the fact that the rate of the entraining AM tone was constant in Experiments 2 and 3, but variable in Experiments 1 and 4.

A regular repetition of a stimulus often leads to a progressive reduction of neural responses to that stimulus. Importantly, such “habituation” or “adaptation” effects are temporally and spectrally specific: Neural activity is suppressed only in response to the expected stimulus, and at the expected moment in time^23,24,26,33^. It is likely that such effects stem from circuits designed to detect novel information, and therefore deviants in both timing and identity^26,27^. In our case, highest sensitivity to novel information would predict a highest number of detected targets in phase and in anti-phase with the preceding AM tone for *f_different_* and *f_same_*, respectively – precisely the opposite as what would be predicted from a tonotopic version of neural entrainment^12,16^.

If a change in stimulus rate is sufficient to “prevent” habituation, then this might explain why we found results to be consistent with neural entrainment only in Experiments 1 and 4 (although some remained inconclusive, see, e.g., *f_different_* in Experiment 4), but not in the other two experiments where rate was constant across trials. Clearly, further experiments – ideally including electrophysiological recordings – are necessary to confirm this opposite relation between neural entrainment and habituation. If confirmed, such a finding would have important consequences for future experimental design, as mixing two counteracting processes might lead to falsely negative or conflicting outcomes. It might also explain why previous results on “rhythmic facilitation of perception and attention”^34^ have not always been unambiguous or straightforward to interpret^30,34–36^ leading to a debate on the concept of entrainment echoes^21,37–39^. Intriguingly, opposite perceptual effects have also been observed in other research fields that examine consequences of stimulus repetition (including “forward masking” and “repetition suppression”) and that have reported both beneficial and detrimental effects on perception or behavior^9,25,26,40–43^.

### Preferred rates for rhythmic entrainment echoes

Preferred rates (*eigenfrequencies*) for auditory neural or perceptual responses have been described before (for review, see ref ^44^). Humans are most sensitive to acoustic spectro-temporal modulations between ~2 and 5 Hz^45^. Neural activity measured with electro- or magnetoencephalography (EEG/MEG) follow such modulations most reliably if they occur in the theta (~4-7 Hz) or gamma (~30-45 Hz) range, but not in-between^46,47^. It has been speculated that these *eigenfrequency* ranges reflect audition’s specialisation to process speech^44,48^, which contains amplitude modulations and changes in linguistic patterns at similar rates^49^. However, most studies have almost exclusively focused on responses during the rhythmic stimulus, when endogenous oscillations are difficult to identify^6^. Indeed, the preferred theta and gamma ranges might simply produce the strongest neural responses because they correspond to stimulus rates that lead to maximally overlapping evoked responses at different cortical levels^45^. It remained, therefore, unclear whether the same preferred rates apply for entrainment echoes, which can be more reliably interpreted as (an “echo” of) entrained endogenous oscillations.

We here identified the preferred rate for rhythmic entrainment echoes as 6 Hz (in Experiment 1, using a wider but coarser range of rates; Fig. 3A) and 8 Hz (in Experiment 4, using a narrower but finer range), respectively. These rates are in line with findings by Ho and colleagues^50^, who found that the onset of broadband noise produces rhythmic fluctuations in auditory perception at similar frequencies (6-8 Hz). Additional research is required to understand why 6-Hz stimulation did not produce a reliable echo in our Experiment 4. Nonetheless, both 6 Hz and 8 Hz are within the dominant AM range of human speech, which - despite a peak at 4-5 Hz - contains significant energy at 6-8 Hz^49,51^. Our findings are therefore compatible with the system’s tuning to process speech. The fact that they are located at the upper limit of this range is in line with the suggestions that *eigenfrequencies* decrease along the auditory hierarchy: As we used tones, not speech, to entrain neural oscillations, perceptual outcomes might have been determined by activity in an earlier cortical stage of cortical processing, where *eigenfrequencies* are likely to be higher than in regions processing more complex linguistic structure^44,45,52^.

We found preferred rates for rhythmic entrainment echoes that differ from those described by Farahbod and colleagues^15^, around 2 Hz. However, in their study only few (3-5) subjects were tested. These subjects were tested extensively, each completing several thousands of trials. This allowed the authors to estimate individual responses very reliably; however, the low number of subjects makes it difficult to draw conclusions on the population level. It is possible that subjects were selected with unusually low preferred rates. How strongly those rates vary on the interindividual level remains to be investigated. Finally, we add that other studies have reported low “natural” rates (1-2 Hz) for audition, but these were often linked to beat perception and audio-motor interactions^53,54^.

### Limitations, open questions and future directions

In all experiments, rhythmic entrainment echoes were relatively short and often restricted to a single cycle. Although this observation is common in the literature^8,9,13^, it does raise some questions. One might wonder whether a rhythmic process that lasts one cycle can be considered a “true” oscillation. This issue is more problematic for standard time-frequency analyses that estimate phase, power, and frequency of the putative oscillation and are therefore more vulnerable to misinterpret other non-oscillatory signals as such. In the current study, however, our analysis of entrainment echoes was guided by clear predictions about both frequency and phase of the hypothesized rhythmic process (Fig. 2). It is relatively unlikely that a non-oscillatory process would have led to best and worst detection in phase and in anti-phase (or vice versa) with the preceding rhythmic stimulus, and intermediate performance in-between. It is also noteworthy that the entrainment echo appeared at delays that are relatively consistent across experiments (~2-3 cycles after the final peak of the AM tone). The absence of an echo in the first cycle after the offset of the rhythmic stimulus (Fig. 3B) can be explained by various effects that are related to this offset (e.g., omission or orientation response^55^). Such effects should be maximal close to stimulus offset and might have masked entrainment echoes, even if it existed, an observation that has been made before^8^.

Independent of this possibility, another factor to be considered is the “usefulness” of echoes for the auditory system, and the consequence for their duration. If these echoes indeed reflect participants’ expectation about stimulus timing, induced by the rhythmicity of the AM stimulus, then it was violated in the current paradigm, as the target was presented at random delays. This would lead to targets randomly coinciding with high- or low-excitability phases of the oscillatory cycle, thus losing the advantage of relevant information being boosted at the high-excitability phase^56^. In such scenarios, it is possible that entrainment echoes are suppressed so that targets are not missed if they occur at the suppressive (low-excitability) phase. The fact that we did observe some rhythmic effects on perception suggests that these echoes were not – or *could not* – be “stopped” immediately after entrainer offset. Nevertheless, it might explain their relatively short duration. This notion – longer or stronger echoes only when they are “useful” for perception (e.g., when most targets are presented at the high-excitability phase of the echo) – needs to be tested in future work. Together, it is likely that target detection is determined by a sophisticated interaction between potentially “automatic” adaptation processes related to stimulus repetition, more higher-level effects of temporal expectation, and attentional mechanisms that have been shown to reverse such expectation effects^57^. Disentangling them is beyond the current study and an exciting endeavour for future work.

## Conclusion

We here demonstrate that the detection of a pure tone target depends on the rhythm of a preceding stimulus if (and only if) the latter is presented between 6 and 8 Hz. Best moments for stimulus detection depended on whether sound frequencies of target and entraining stimulus match, supporting the notion of tonotopy in rhythmic entrainment echoes. Nevertheless, these echoes seem more complex than those predicted from initial theories of neural entrainment. This complexity might be partly due to the fact that neural entrainment and repetition-related adaption exercise competing, opposite influences on perception.

## Acknowledgements

The authors thank Andrea Alamia, Florian Kasten and Jules Erkens for fruitful discussions.

## Supplementary Figures

**Figure S1.**
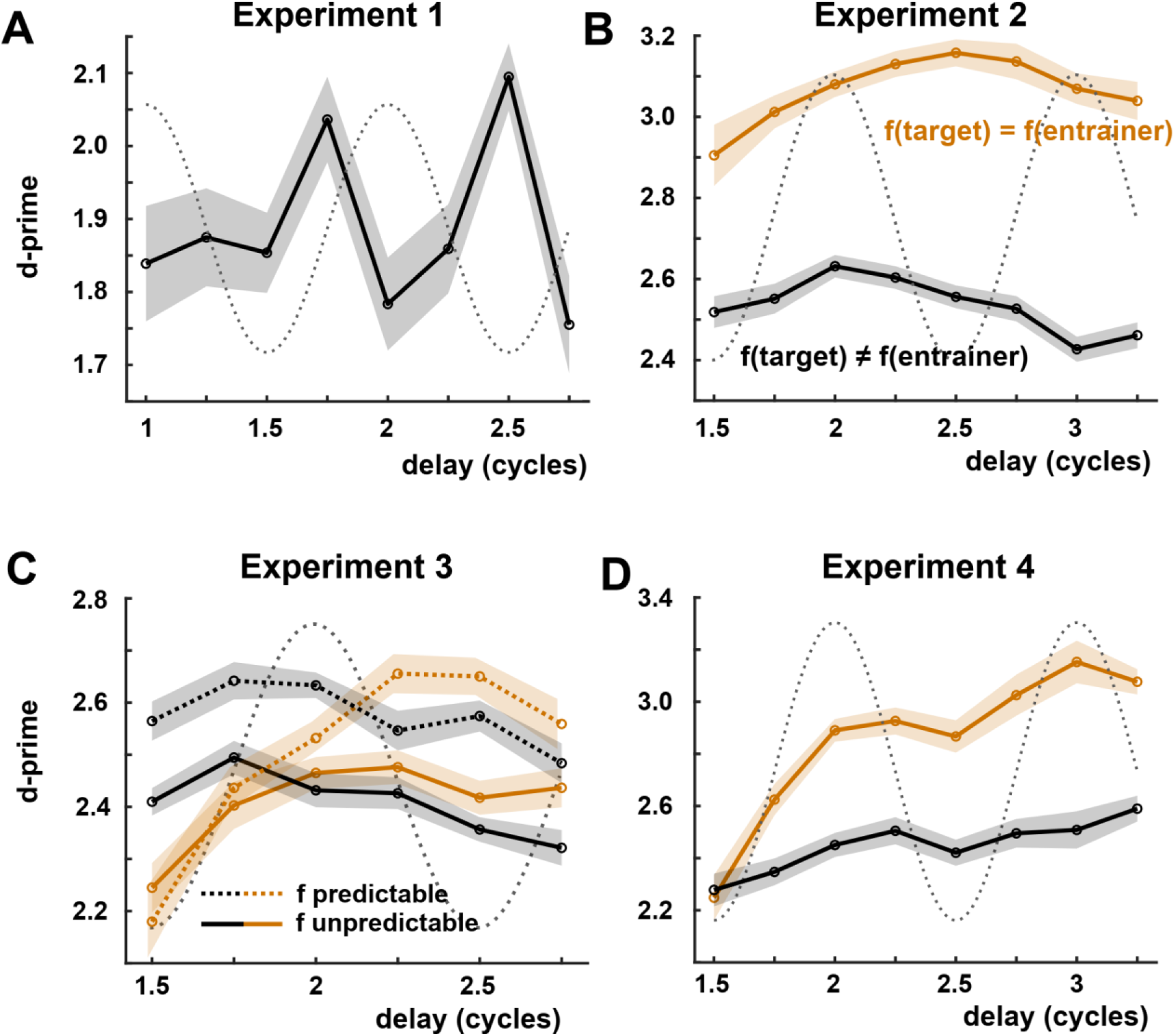
D-prime as a function of target delay relative to the rhythmic AM tone. Same conventions as for Fig. B3. Panels correspond to Figs. 3B, 4A, 5A,B, and 6B, which show similar fluctuations in the proportion of detected targets, respectively.

## References

1. Lakatos, P., Karmos, G., Mehta, A. D., Ulbert, I. & Schroeder, C. E. Entrainment of neuronal oscillations as a mechanism of attentional selection. Science 320, 110–113 (2008).

2. Walter, V. J. & Walter, W. G. The central effects of rhythmic sensory stimulation. Electroencephalogr. Clin. Neurophysiol. 1, 57–86 (1949).

3. Fröhlich, F. & McCormick, D. A. Endogenous Electric Fields May Guide Neocortical Network Activity. Neuron 67, 129–143 (2010).

4. Obleser, J. & Kayser, C. Neural Entrainment and Attentional Selection in the Listening Brain. Trends Cogn. Sci. 23, 913–926 (2019).

5. Lakatos, P., Gross, J. & Thut, G. A New Unifying Account of the Roles of Neuronal Entrainment. Curr. Biol. 29, R890–R905 (2019).

6. Zoefel, B., ten Oever, S. & Sack, A. T. The Involvement of Endogenous Neural Oscillations in the Processing of Rhythmic Input: More Than a Regular Repetition of Evoked Neural Responses. Front. Neurosci. 12, (2018).

7. Keitel, C., Quigley, C. & Ruhnau, P. Stimulus-Driven Brain Oscillations in the Alpha Range: Entrainment of Intrinsic Rhythms or Frequency-Following Response? J. Neurosci. 34, 10137–10140 (2014).

8. Bree, S. van, Sohoglu, E., Davis, M. H. & Zoefel, B. Sustained neural rhythms reveal endogenous oscillations supporting speech perception. PLOS Biol. 19, e3001142 (2021).

9. Saberi, K. & Hickok, G. Forward entrainment: Psychophysics, neural correlates, and function. Psychon. Bull. Rev. (2022) doi:10.3758/s13423-022-02220-y.

10. Spaak, E., Lange, F. P. de & Jensen, O. Local Entrainment of Alpha Oscillations by Visual Stimuli Causes Cyclic Modulation of Perception. J. Neurosci. 34, 3536–3544 (2014).

11. Kösem, A. et al. Neural Entrainment Determines the Words We Hear. Curr. Biol. 28, 2867–2875.e3 (2018).

12. Lakatos, P. et al. The spectrotemporal filter mechanism of auditory selective attention. Neuron 77, 750–761 (2013).

13. Hickok, G., Farahbod, H. & Saberi, K. The Rhythm of Perception: Entrainment to Acoustic Rhythms Induces Subsequent Perceptual Oscillation. Psychol. Sci. 26, 1006–1013 (2015).

14. Fröhlich, F. Chapter 3 - Experiments and models of cortical oscillations as a target for noninvasive brain stimulation. in Progress in Brain Research (ed. Bestmann, S.) vol. 222 41–73 (Elsevier, 2015).

15. Farahbod, H., Saberi, K. & Hickok, G. The rhythm of attention: Perceptual modulation via rhythmic entrainment is lowpass and attention mediated. Atten. Percept. Psychophys. 82, 3558–3570 (2020).

16. O’Connell, M. N., Barczak, A., Schroeder, C. E. & Lakatos, P. Layer Specific Sharpening of Frequency Tuning by Selective Attention in Primary Auditory Cortex. J. Neurosci. 34, 16496–16508 (2014).

17. Peer, E., Samat, S., Brandimarte, L. & Acquisti, A. Beyond the Turk: Alternative Platforms for Crowdsourcing Behavioral Research. https://papers.ssrn.com/abstract=2594183 (2016).

18. de Leeuw, J. R. jsPsych: A JavaScript library for creating behavioral experiments in a Web browser. Behav. Res. Methods 47, 1–12 (2015).

19. Woods, K. J. P., Siegel, M. H., Traer, J. & McDermott, J. H. Headphone screening to facilitate web-based auditory experiments. Atten. Percept. Psychophys. 79, 2064–2072 (2017).

20. Zoefel, B., Gilbert, R. A. & Davis, M. H. Intelligibility improves perception of timing changes in speech. BiorXiv 2022.05.18.492430 Preprint at https://doi.org/10.1101/2022.05.18.492430 (2022).

21. Saberi, K. & Hickok, G. Confirming an antiphasic bicyclic pattern of forward entrainment in signal detection: A reanalysis of Sun et al. (2021). Eur. J. Neurosci. 56, 5274–5286 (2022).

22. Schroeder, C. E. & Lakatos, P. Low-frequency neuronal oscillations as instruments of sensory selection. Trends Neurosci. 32, 9–18 (2009).

23. Herrmann, B., Henry, M. J. & Obleser, J. Frequency-specific adaptation in human auditory cortex depends on the spectral variance in the acoustic stimulation. J. Neurophysiol. 109, 2086–2096 (2013).

24. Costa-Faidella, J., Baldeweg, T., Grimm, S. & Escera, C. Interactions between “What” and “When” in the Auditory System: Temporal Predictability Enhances Repetition Suppression. J. Neurosci. 31, 18590–18597 (2011).

25. Lange, K. Brain correlates of early auditory processing are attenuated by expectations for time and pitch. Brain Cogn. 69, 127–137 (2009).

26. Ulanovsky, N., Las, L. & Nelken, I. Processing of low-probability sounds by cortical neurons. Nat. Neurosci. 6, 391–398 (2003).

27. Khouri, L. & Nelken, I. Detecting the unexpected. Curr. Opin. Neurobiol. 35, 142–147 (2015).

28. Hanslmayr, S., Matuschek, J. & Fellner, M.-C. Entrainment of prefrontal beta oscillations induces an endogenous echo and impairs memory formation. Curr. Biol. 24, 904–909 (2014).

29. Large, E. W. & Jones, M. R. The dynamics of attending: How people track time-varying events. Psychol. Rev. 106, 119–159 (1999).

30. Bauer, A.-K. R., Jaeger, M., Thorne, J. D., Bendixen, A. & Debener, S. The auditory dynamic attending theory revisited: A closer look at the pitch comparison task. Brain Res. 1626, 198–210 (2015).

31. Jones, M. R., Moynihan, H., MacKenzie, N. & Puente, J. Temporal aspects of stimulus-driven attending in dynamic arrays. Psychol. Sci. 13, 313–319 (2002).

32. Graaf, T. A. de et al. Alpha-Band Rhythms in Visual Task Performance: Phase-Locking by Rhythmic Sensory Stimulation. PLOS ONE 8, e60035 (2013).

33. Micheyl, C., Tian, B., Carlyon, R. P. & Rauschecker, J. P. Perceptual Organization of Tone Sequences in the Auditory Cortex of Awake Macaques. Neuron 48, 139–148 (2005).

34. Haegens, S. & Zion Golumbic, E. Rhythmic facilitation of sensory processing: A critical review. Neurosci. Biobehav. Rev. 86, 150–165 (2018).

35. Vilà-Balló, A., Marti-Marca, A., Torralba Cuello, M., Soto-Faraco, S. & Pozo-Rosich, P. The influence of temporal unpredictability on the electrophysiological mechanisms of neural entrainment. Psychophysiology 59, e14108 (2022).

36. Barne, L. C., Cravo, A. M., de Lange, F. P. & Spaak, E. Temporal prediction elicits rhythmic preactivation of relevant sensory cortices. Eur. J. Neurosci. 55, 3324–3339 (2022).

37. Sun, Y., Michalareas, G. & Poeppel, D. The impact of phase entrainment on auditory detection is highly variable: Revisiting a key finding. Eur. J. Neurosci. 55, 3373–3390 (2022).

38. Lin, W. M. et al. No behavioural evidence for rhythmic facilitation of perceptual discrimination. Eur. J. Neurosci. 55, 3352–3364 (2022).

39. Saberi, K. & Hickok, G. A critical analysis of Lin et al.’s (2021) failure to observe forward entrainment in pitch discrimination. Eur. J. Neurosci. 56, 5191–5200 (2022).

40. Todorovic, A. & Lange, F. P. de. Repetition Suppression and Expectation Suppression Are Dissociable in Time in Early Auditory Evoked Fields. J. Neurosci. 32, 13389–13395 (2012).

41. Segaert, K., Weber, K., de Lange, F. P., Petersson, K. M. & Hagoort, P. The suppression of repetition enhancement: A review of fMRI studies. Neuropsychologia 51, 59–66 (2013).

42. Jesteadt, W., Bacon, S. P. & Lehman, J. R. Forward masking as a function of frequency, masker level, and signal delay. J. Acoust. Soc. Am. 71, 950–962 (1982).

43. Sohoglu, E. & Chait, M. Detecting and representing predictable structure during auditory scene analysis. eLife 5, e19113 (2016).

44. Zoefel, B. & Kösem, A. Neural Dynamics: Speech is Special. PsyArXiv Preprint at https://doi.org/10.31234/osf.io/eyrtb (2022).

45. Edwards, E. & Chang, E. F. Syllabic (~2–5 Hz) and fluctuation (~1–10 Hz) ranges in speech and auditory processing. Hear. Res. 305, 113–134 (2013).

46. Teng, X., Tian, X., Rowland, J. & Poeppel, D. Concurrent temporal channels for auditory processing: Oscillatory neural entrainment reveals segregation of function at different scales. PLOS Biol. 15, e2000812 (2017).

47. Teng, X. & Poeppel, D. Theta and Gamma Bands Encode Acoustic Dynamics over Wide-Ranging Timescales. Cereb. Cortex 30, 2600–2614 (2020).

48. Poeppel, D. & Assaneo, M. F. Speech rhythms and their neural foundations. Nat. Rev. Neurosci. 21, 322–334 (2020).

49. Ding, N. et al. Temporal modulations in speech and music. Neurosci. Biobehav. Rev. 81, 181–187 (2017).

50. Ho, H. T., Leung, J., Burr, D. C., Alais, D. & Morrone, M. C. Auditory Sensitivity and Decision Criteria Oscillate at Different Frequencies Separately for the Two Ears. Curr. Biol. 27, 3643–3649.e3 (2017).

51. Varnet, L., Ortiz-Barajas, M. C., Erra, R. G., Gervain, J. & Lorenzi, C. A cross-linguistic study of speech modulation spectra. J. Acoust. Soc. Am. 142, 1976 (2017).

52. Giraud, A.-L. et al. Representation of the Temporal Envelope of Sounds in the Human Brain. J. Neurophysiol. 84, 1588–1598 (2000).

53. Weineck, K., Wen, O. X. & Henry, M. J. Neural synchronization is strongest to the spectral flux of slow music and depends on familiarity and beat salience. eLife 11, e75515 (2022).

54. Zalta, A., Petkoski, S. & Morillon, B. Natural rhythms of periodic temporal attention. Nat. Commun. 11, 1051 (2020).

55. Hughes, H. C. et al. Responses of Human Auditory Association Cortex to the Omission of an Expected Acoustic Event. NeuroImage 13, 1073–1089 (2001).

56. Zoefel, B. & VanRullen, R. Oscillatory Mechanisms of Stimulus Processing and Selection in the Visual and Auditory Systems: State-of-the-Art, Speculations and Suggestions. Front. Neurosci. 11, (2017).

57. Kok, P., Rahnev, D., Jehee, J. F. M., Lau, H. C. & de Lange, F. P. Attention reverses the effect of prediction in silencing sensory signals. Cereb. Cortex 22, 2197–2206 (2012).

